# Synchrony surfacing: epicortical recording of correlated action potentials

**DOI:** 10.1101/329342

**Authors:** Tobias Bockhorst, Florian Pieper, Gerhard Engler, Thomas Stieglitz, Edgar Galindo-Leon, Andreas K. Engel

**Affiliations:** Department of Neurophysiology and Pathophysiology, University Medical Center Hamburg-Eppendorf, Hamburg, Germany; Department of Microsystems Engineering, University of Freiburg, Freiburg, Germany

**Keywords:** Electrocorticography, synchronous spiking, assembly-coding, visual cortex, auditory cortex, brain-machine interfaces

## Abstract

Synchronous spiking of multiple neurons is a key phenomenon in normal brain function and pathologies. Recently, approaches to record spikes from the intact cortical surface using small high-density arrays of microelectrodes have been reported. It remained unaddressed how epicortical spiking relates to intracortical unit activity. We introduce a mesoscale approach using an array of 64 electrodes with intermediate diameter (250 µm) and combined large-coverage epicortical recordings in ferrets with intracortical recordings via laminar probes. Empirical data and modeling strongly suggest that our epicortical electrodes selectively captured synchronized spiking of neurons in the subjacent cortex. As a result, responses to sensory stimulation were more robust and less noisy as compared to intracortical activity, and receptive field properties were well preserved in epicortical recordings. This should promote insights into assembly-coding beyond the informative value of subdural EEG or single-unit spiking, and be advantageous to real-time applications in brain-machine interfacing.

**Significance statement:** Electrocorticography allows chronic, low-noise recordings from the intact cortical surface - a prerequisite for investigations into brain network dynamics and brain-machine interfaces. Novel electrodes can capture spiking activity at the surface, which should boost precision in the spatial - and time domain, compared to conventional EEG-like measurements. To clarify how surface spiking relates to intracortically fired action potentials, we recorded both types of signal simultaneously from sensory cortices in anesthetized ferrets. Results suggest that mesoscale (250 µm) surface electrodes can selectively capture synchronized spiking from nearby cortical columns, which reduces contamination by non-representative, jittering spikes. Given the high relevance of neural synchrony for sensorimotor and cognitive processing, the novel methodology may improve signal decoding in brain-machine interface approaches.

**Author contributions:** E.G.L., T.B. and A.K.E. conceptualized the research; E.G.L. and F.P. performed experiments; T.B. and E.G.L. wrote Matlab routines for data analysis; T.B. and E.G.L. analyzed the data; T.S. provided technical resources; T.B., E.G.L. and A.K.E. wrote the manuscript; G.E. administrated the project; A.K.E. acquired funding.

## Introduction

Recordings with electrocorticographic (ECoG) arrays allow prolonged, spatially resolved measurements of activity in distributed cortical networks with low noise and minimal invasiveness [1,2,3]. They are also a key to map cortical functions and to study cognitive and motor processes in the healthy brain [4]. In clinical applications they serve to localize epileptogenic zones prior to surgery [5,6]. In recent years, arrays with a large number of small-sized electrodes (high-density ECoG arrays) have been developed and successfully used in brain machine interfaces (BMI), for instance, in the decoding of natural grasp types [7] and word recognition [8,9] in humans. Most electrode arrays were designed to cover small cortical subregions of interest with high spatial resolution. Yet, such local high-density ECoG arrays are not well suited to study brain dynamics at large-scale with concurrent recordings from multiple cortical areas.

Some studies have implied that an analysis of spiking activity would add relevant information to ECoG recordings [10]. In particular, in the decoding of grasp types, accuracy was highest for activity in the high gamma band, a presumed correlate of coordinated assembly spiking [11,7]. Measurements of spiking activity in response to stimulation or task demands, are considered the gold standard for studies on neural coding [12]. Recent reports have suggested that action potentials, possibly from individual neural units, can be recorded at the cortical surface using arrays of microscale electrodes [13]. The significance of the obtained signal in processes such as stimulus coding, decision making or motor control remains a matter of debate. Moreover, it remains to be elucidated how epicortically measured spiking relates to its intracortical substrate, in particular with respect to the coding of stimulus properties and the correlation structure of spiking activity in the subjacent cortical columns.

In the present study, we demonstrate that an ECoG-array of mesoscale geometry (250 µm diameter, 1.5 mm spacing) is suited to capture representative assembly spiking at macrocolumn resolution, while being small enough to prevent the blurring of tuning properties in visual and auditory regions. Its principle design is very adaptable, allowing to cover large part of a hemisphere of a monkey brain [14,15]. Former studies based on this design have focused on the frequency range of local field potentials (<300 Hz), thus capturing subthreshold activity. We now explore its suitability for spike recordings by comparing the thresholded high-frequency component of both, ECoG and intracortical signals.

To this end, we employed a combination of ECoG and intracortical laminar probe recordings to simultaneously obtain both types of signal. Average waveform features of epicortical spikes still resembled those of presumed single-unit action potentials recorded from the cortical surface with micro-scale arrays [13]. We show that the epicortically measured multiunit spiking correlates with intracortical multiunit activity both during ongoing activity and under sensory stimulation. Importantly, we demonstrate that epicortical multiunit signals are best predicted from intracortical recordings when taking only synchronized spikes into account. As a result, epicortical responses were stronger as well as less variable in terms of latency, amplitude and duration. Furthermore, the coding of stimulus properties in the auditory and visual domain was well preserved; in fact, visual receptive fields were more confined in the epicortical data. We propose that, with the mesoscale approach employed here, epicortically measured responses reflect mainly synchronized spikes from intracortical neural assemblies.

## Materials and methods

Data presented in this study were collected from 4 adult female ferrets (*Mustela putorius*, Euroferret, Dybbølsgade, Denmark). All experiments were approved by the Hamburg state authority for animal welfare (BUG-Hamburg) and were performed in accordance with the guidelines of the German Animal Protection Law.

### Custom ECoG design

Recordings were carried out using a custom-made ECoG array that matches the anatomy of the left hemisphere of the ferret brain (Fig 1A). The array consists of 64 platinum electrodes with a diameter of 250 μm each, embedded in a thin-film polyimide foil of 10 µm thickness and arranged in a hexagonal formation with an interelectrode distance of 1.5 mm. For details, see [14].

### Surgery

Details on the surgical procedure have been reported earlier [2]. Briefly, animals were initially anesthetized with an injection of ketamine (15 mg/kg) and medetomidine (0.08 mg/kg). A glass tube was inserted into the trachea to allow artificial ventilation of the animal and to supply isoflurane (approx. 0.4%; exact concentration adjusted as to avoid burst-suppresion states) for maintenance of anesthesia. Body temperature, heart rate and end-tidal CO_2_ concentration were constantly monitored throughout the whole experiments to maintain the state of the animal. To prevent dehydration, a cannula was inserted into the femoral vein to deliver a continuous infusion of 0.9% NaCl, 0.5 % NaHCO_3_ and pancuronium bromide (6 µg/kg/h). To ensure monocular stimulation, the left eye was occluded. Finally, to avoid desiccation of the cornea, a contact lens was placed on the right eye.

The temporalis muscle was folded back, such that a craniotomy could be performed over the left posterior cortex. The dura was carefully removed before placing the ECoG array on the surface of the cortex. In each animal, the position of the array on the cortex was photographed with a Zeiss OPMI pico microscope to obtain the exact location of ECoG electrodes. The position of ECoG electrode grids was projected onto a scaled model ferret brain that contained a map of the functional cortical areas (20 in total) [16].

Once the ECoG array was fully placed, two silicon probes of 32 electrodes linearly distributed (A1×32-15mm-50-177, NeuroNexus Technologies) were introduced through holes in the ECoG array and manually advanced via a micromanipulator (David Kopf Instruments) under visual inspection through the microscope. One of the probes was introduced in an early visual area, the second in an early auditory area (Figs 1A, 2A). The probes were advanced gently just until the uppermost electrode caught a physiological signal, indicating that it had just entered the cortex. During the whole experimental session, saline solution was gently dropped upon the preparation to prevent exsiccation.

### Sensory stimulation

To ensure controlled conditions for sensory stimulation, all experiments were carried out in a dark, sound-attenuated anechoic chamber (Acoustair, Moerkapelle, Netherlands). Auditory and visual stimuli were generated using the Psychophysics Toolbox [17] in Matlab (Mathworks Inc, Natick, MA). Acoustic stimuli were digitalized at 96kHz and delivered with an RME soundcard (RME HDSPe AIO Intelligent Audio Solutions) through a Beyerdynamic T1 speaker located 15 cm from the animal’s right ear. Prior to stimulation, the sound delivery system was calibrated using a Brüel and Kjær (B&K, Nærum, Denmark) free-field 4939 microphone coupled to a B&K 2670 pre-amplifier and 2690 amplifier. Visual stimuli were presented on an LCD monitor (Samsung SyncMaster 2233, frame rate 100 Hz) placed 28 cm in front of the animal to cover a visual field of 52°x52°.

*Visual Stimulation*: Visual stimuli were presented monocularly through the right eye. For mapping of receptive fields, we presented small-field flashes (circular patches of 2° in diameter with Gaussian amplitude distribution) of 10 ms duration distributed over a 17×17 grid. The spatial position of the flash was randomly selected and the time between stimuli varied randomly between 150 and 200 ms. In total, each position was probed ten times.

In addition, to measure feature selectivity, visual stimulation was carried out using drifting Gabor patches with an extent of 14 degrees (full width at half maximum), a spatial frequency of 2 cycles/deg and a temporal frequency of 5 deg/sec, drifting in 4 directions (0°, 90°, 180° and 270°). Each stimulus was presented 10 times.

*Auditory stimulation*: Spectro-temporal receptive fields (tuning curves) were derived at 65 dB SPL by presenting a set of 38 tones with logarithmically spaced frequencies ranging from 400 Hz to 32 kHz (200 ms duration, including 10 ms cos^2^ ramps at onset and offset). Frequencies were presented 10 times in a semi-randomized manner (first all frequencies were presented before moving to the next repetition).

### Data acquisition and analysis

To record multiunit activity from the ECoG and the intracortical laminar probe, signals were band-pass filtered between 0.5 Hz to 8 kHz, digitized at 44 kHz and sampled with an AlphaLab SnR^TM^ recording system (Alpha Omega Engineering, Nazareth, Israel). All analyses of neural data presented in this study were performed offline after the completion of experiments using custom-written Matlab scripts (Mathworks). For spike detection, both ECoG and intracortical signals were band-pass filtered between 300 Hz and 5 kHz, transformed to zero median and normalized by the long-tail-corrected standard deviation (median of the absolute values divided by 0.6745; according to [18] of the time series. Spike times were identified as local minima below a threshold of -3 standard deviations (artifact rejection threshold: -20 standard deviations), with no refractoriness assumed for the multi-unit activity measured here.

For each experiment, we calculated the degree to which spiking activity (spike trains convolved with 5 ms boxcar-kernel) at the 32 intracortical electrodes along the laminar probe was correlated with any of the 64 epicortical electrodes. The resultant 32×64 matrix of correlation strengths was then screened to identify the pair of electrodes that yielded the best correlation. These best-correlated pairs provided the data underlying comparisons within pairs of epi- and intracortical electrodes.

In order to quantify the tuning properties of the visual recordings, we defined the modulation index MI as 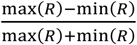 with *R* being the vector of response strengths to different movement directions. The same index was used for quantification of general response strength and for quantifying the asymmetry of spike-waveforms.

### Modeling

We developed a forward model to explain the surface activity based on the intracortical spiking activity. For modeling of receptive fields, we first selected those units whose spatial response patterns resembled the principle structure of a receptive field (response > 3 STD for at least 9 consecutive locations on the visual grid). A sliding time-window then detected spikes that occurred in concert across contacts of the intracortical probe within a synchrony window that varied between simulations, ranging from 0.5 to 100 ms. Those spikes were combined into a single spike train on which a standard analysis of mapping responses was performed to estimate receptive field diameters at half-maximum response amplitude. Spikes that occurred in exact simultaneity (zero delay) across more than one recording channel were considered as resulting from the same neuron. We also investigated the effect of spike waveform on the sum of extracellular potentials. For this purpose, we converted the original spike trains (of the selected channels) into a binary continuous signals of extracellular potentials (sampling frequency 44 kHz) by convolution with a spike-waveform kernel (regular and fast spikes, Fig 2G) [19]. Finally, we summed the resultant extracellular potentials and analyzed the resultant stimulus response. A similar procedure was applied to model responses to sensory stimuli, by convolving binary trains of synchronous spikes with a boxcar kernel (10 ms). In addition, effects of thresholding the assembly size, i.e., the minimum number of coincidences with other units required to include a spike (see Fig 3) were explored.

## Results

Data were obtained from a total of 4 adult female ferrets (*Mustela putorius*) under isoflurane anesthesia. In each animal, data were recorded across a time span of 10-12 hours, thus balancing out any possible confounds resulting from slight changes in brain state related to circadian epochs or effective levels of anesthesia. Epicortical multiunit activity was recorded using custom-made ECoG-arrays with 64 electrodes covering the posterior half of the left hemisphere of the ferret brain (Fig 1A). Laminar probes with 32 contacts were inserted in primary visual and auditory cortices to capture intracortical spike activity concurrently to epicortical recordings (Fig 1A).

**Fig 1.**
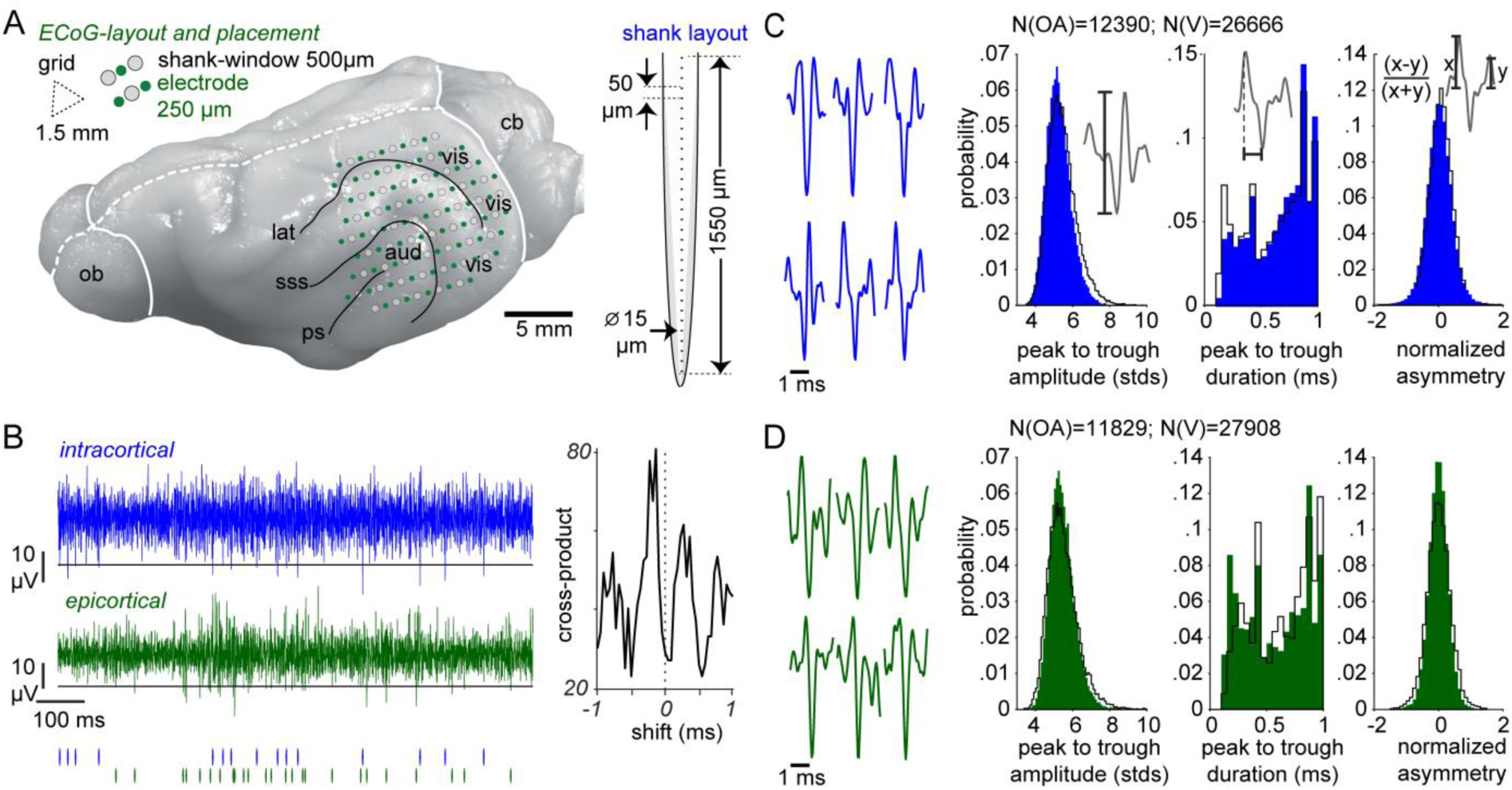
Recording approach and basic features of epi- and intracortical spikes. (A) (Left) Hexagonal geometry of the ECoG-array and its position on the posterior half of the left hemisphere, including coverage of primary visual (vis) and auditory (aud) areas. Black contours: sulci in the regions covered (lat: lateral, sss: suprasylvian, ps: pseudosylvian); dashed white line: interhemispheric fissure; cb: cerebellum; ob: olfactory bulb. (Right) Geometry of the single-shank linear probe (NeuroNexus) inserted through the shank-windows in the ECoG-array. (B) Example traces (300-5000 Hz; 1s activity) of the intracerebral (blue, upper trace) and epicortical (green, lower trace) signal. Data are based on the best-correlated electrode pair from a randomly chosen measurement. Black horizontal lines: spike detection thresholds, corresponding to three standard deviations of the noise. Spike rasters and the cross-correlogram (cross-product of binary spike trains; calculated for the entire measurement duration of 25 min) depicted below show that the signals are not trivially associated by crosstalk, which would be indicated by zero-lag spike cross-correlation. (C and D) Representative examples of detected waveforms and histograms of major waveform-features, including peak-to-trough amplitude (in unit standard deviations), peak-to-trough duration and normalized asymmetry, for intracortical and epicortical spike recordings. Data are from one experimental animal (cf. S1 Fig for data from the other animals). Coloured bars indicate data from visual stimulation (V), overlaid black contours indicate data from ongoing activity (OA). N, number of spikes from OA and V. Note that distributions of spike-waveform features are very similar for both sources and effectively identical for ongoing and visually elicited activity. Moreover, distributions of peak-to-trough durations are bimodal for both intracortical and epicortical data.

### Basic features of epicortical and intracortical spikes are highly similar

We first compared basic features of the spikes detected at intracortical and epicortical electrode contacts. For each of the experiments, this comparison was made for those ECoG contacts and intracortical channels that showed the highest correlation of the spike density time series. Spikes were detected both in ongoing activity and under sensory stimulation. We observed no trivial, i.e., instantaneous correlation between exact spike times at best-correlated pairs of intra- and epicortical electrodes (Fig 1B; S1 Fig B, D, F), which rules out any substantial contribution of correlated high-frequency electrical noise. Principle waveform features were remarkably similar for intracortical and epicortical spikes (Fig 1C, D). The distributions of peak-to-trough durations were often bimodal in both signals (Fig 1C, D; S1 Fig A, B, C), which might suggest that the contribution of fast - and slow spiking cells (i.e., inhibitory interneurons and excitatory neurons) was still visible in the epicortical recordings.

### Epicortical responses show similar tuning but smaller receptive fields than intracortical recordings

As a next step, we investigated responses to different types of sensory stimuli as well as the mapping of intracortical response properties to epicortical activity. We compared multiunit signals from all ECoG contacts and all intracortical channels that showed a receptive field. If the transfer from intracortical to epicortical spike rates involves spatial averaging (as expected for the size of our ECoG electrode contacts) or nonlinearities (as might result from intracortical spike synchrony) this might affect the degree to which information on sensory stimuli may be preserved or diminished in epicortical signals. To address this, we investigated whether receptive field size and direction tuning in early visual areas, as well as frequency tuning in primary auditory cortex were preserved in the epicortical data.

If retinotopy was blurred by superposition of spatially distinct receptive fields, the resulting epicortical receptive fields should be substantially broader. Interestingly, we found that receptive fields of epicortical multiunit activity were significantly smaller in diameter (full width at half maximum) compared to the intracortical receptive fields (N_intra_= 78, N_epi_= 72, Wilcoxon test, p< 0.0001; Fig 2A,B,D). To quantify the strength of directional tuning, response contrasts for gratings presented in four movement directions were calculated. For each recording location we calculated a mean modulation-index, reflecting the contrast between strongest and weakest responses, as a measure of directional sensitivity. Across populations of epi- and intracortical recordings we found no significant difference between both groups (Fig 2D).

Fig 2C juxtaposes patterns of tonotopical (pure tone frequency) tuning in epi-vs. intracortical measurements. Similar to visual receptive fields, differences in auditory tuning between epicortical and intracortical spiking should reflect the degree of blur in the epicortical signal. Tuning widths (in octaves) in the intracortical data tended to be slightly narrower but the difference did not reach statistical significance. The auditory tuning curves mostly resembled their intracortical counterparts in shape, but observations of double peaks in epicortical tuning profiles suggest that occasional superposition of two or more single-peak curves could explain the trend toward higher half-maximum bandwidths on average (Fig 2A,C,D).

**Fig 2.**
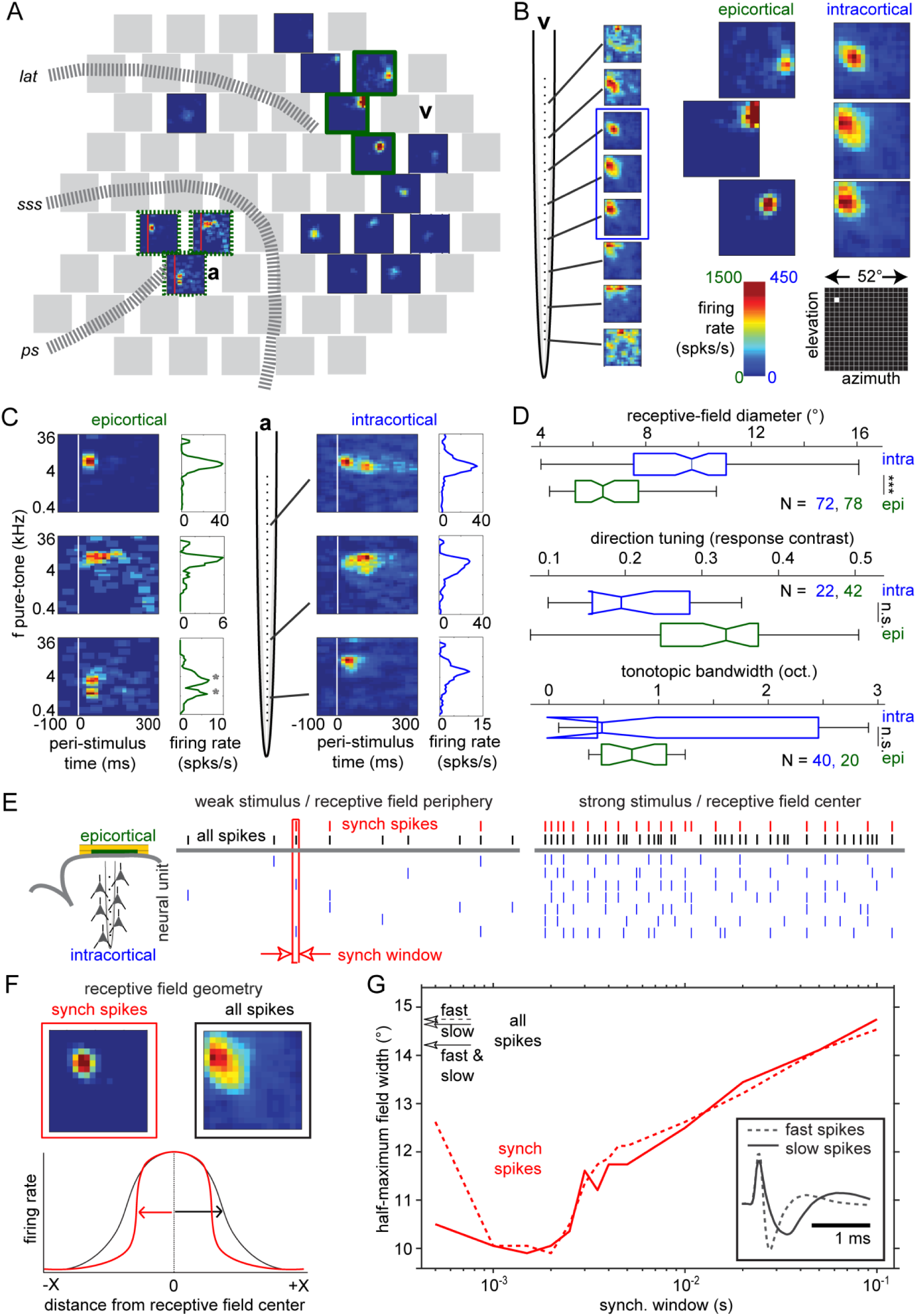
Relation between intracortical and epicortical spiking: receptive fields and tuning to stimulus features. Comparisons include spatial (visual) and spectro-temporal (auditory) receptive fields as well as tuning to orientation of visual stimuli. (A-D) Experimental data; (E-G) modeling. (A-C) Example data from one experimental animal. (A) Topography of visual and auditory responses, indicated as receptive field maps plotted at the position of the responsive contacts. a, v: positions of silicon probes for intracortical recording of auditory and visual activity, respectively. (B) Visual receptive fields defined by responses to small flashes presented in a mapping grid of 17×17 positions (52°x52°). (Left) Receptive fields measured at different electrodes along the intracortical probe. (Right) Close-up comparisons of receptive fields from epicortical and intracortical recordings (obtained at the sites framed in blue on the left, and in green in panel A, respectively). (C) Spectro-temporal receptive fields and frequency tuning-curves obtained from responses to auditory pure-tone stimulation. Asterisks in one plot of epicortical tuning indicate a double-peaked curve. (D) Population analysis of response properties for intracortical (blue) and epicortical (green) recordings. N, number of recordings. (Top) Receptive field size, measured as full-width at half of the maximum (*** p<10^−5^). (Middle) Tuning to movement direction of drifting Gabor patterns (p=0.06). (Bottom) Bandwidth of auditory tuning curves, measured as full-width at half of the maximum and expressed in octaves (p>0.05). (E) Schematic illustration of the hypothesis that the ECoG contacts preferentially pick up synchronized spikes. The intracortical probe records the spiking activity of multiple cells through different electrode channels (black). We assume that only spikes that are coincident within a narrow time window (‘synch window’) are detected by the epicortical contact (green). (Left) For weak stimulation or stimulation in the receptive field periphery, only a small proportion of the spikes will be synchronized (red spikes). (Right) For optimal stimulation, the probability of synchronized spikes (red) in the local cortical columns strongly increases. This may lead to a sharpening of receptive fields as shown in F. (F) Examples of receptive fields obtained taking only synchronized spikes (top left) or all spikes (top right). Firing rates are normalized to maximum response in each case. (G) Size of receptive fields (full width at half-maximum) obtained from all spikes (arrows on vertical axis) versus synchronized spikes (red graphs), respectively. For synchronized spikes, field width is plotted as a function of the width of the synchrony window applied. Each spike train was convolved either with a kernel of fast spikes (dotted line) and regular spikes (solid lines).

### Epicortical receptive fields likely result from synchronized intracortical spikes

We implemented a forward model (Fig 2E-G) to assess whether selective propagation of synchronous intracortical spikes may explain the abovementioned size reduction of receptive fields in the epicortical data. We hypothesized that the activity projected to the surface is a thresholded sum of intracortical activities aligned within a synchrony window σ. The schematic diagram in Fig 2E illustrates how spiking activity projects to the cortical surface at the periphery vs. center of the receptive field, if filtering for synchronous spikes (red spike train) is assumed. We tested this hypothesis for the recorded data, varying σ between 0.5 and 100 ms, and found that the size of the epicortical receptive fields considerably dropped for values of σ in the range of 1-2 ms (Fig 2G). The reduction in receptive field size was about 30% of the reference value obtained under linear summation including all intracortical spikes. This relation matches the observed relation of receptive field sizes between the epicortical vs. intracortical data (Fig 2D). Spike waveform did not affect this result substantially, suggesting that the proposed mechanism applies in the same way to any type of neuron.

### Strength and variability of epicortical responses are best explained by synchrony

As a next step, we compared the response dynamics for epicortical and intracortical responses. If epicortical signals reflect mainly synchronized spiking, we would expect responses to be less variable since noisy components that are not synchronized across units and strongly variable across trials might get suppressed. Depending on the degree of synchrony as well as the number of units picked up by the ECoG contact, epicortical responses may also be higher in amplitude than single-contact intracortical responses.

Average spike counts for ongoing activity were only slightly higher on average, but more variable, in epicortical measurements (Fig 3C). By contrast, epicortical responses to visual stimuli (full-field flashes, 10ms) were markedly higher in amplitude than intracortical responses (Fig 3A,B,D). When trial-averaged responses were compared by means of computing the bin-wise ratio of peri-stimulus time histograms, epicortical rates were found to exceed intracortical ones significantly (p<<0.0001; Fig 3A), being about three times as high on average. Despite the observed variability in ongoing activity, a significant, strong and consistent increase was also observed for modulation indices, i.e., relative amplitudes of epicortical responses that are normalized to the trial-resolved level of pre-stimulus activity (p<<0.001; Fig 3D). Trial-level statistics (S1 Table) confirmed that epicortical responses were not only higher in amplitude than intracortical single-contact responses, but had comparable or smaller latencies, comparable or higher durations and less variability in each of these features.

To test whether epicortical response dynamics might be best explained by spike synchrony, the same forward model applied to receptive fields was used (Fig. 3 E-G). To model the time course of epicortical responses, the rasters of the 32 spike trains recorded by the contacts of the intracortical probe were screened for coincidences of spikes. We applied a bin width of 2ms suggested as the relevant synchrony window in the modeling of receptive field diameters described above. We then tested the effects of including variable minimum numbers of coincident spikes recorded across contacts of the intracortical probe, thus simulating a variation in the size of an assembly of synchronized cells. The number of required coincidences per 2ms time bin was varied between 1 (all spikes accepted) and 10 (high degree of synchrony). To produce peri-stimulus-time histograms, the trains of accepted synchronous spikes were convolved with a 10 ms boxcar kernel. In general, an increase in the number of required coincidences resulted in decreased average rate and increased fluctuation (quantified in terms of the rate’s coefficient of variation) of pre-stimulus spiking (Fig 3E), while relative response strength (modulation indices, reflecting response strength above pre-stimulus activity) increased and stabilized (Fig 3F). These effects saturated (mean pre-stimulus rate, relative response strength), respectively increased (variability), when level of about 7 coincidences per bin was exceeded. Further analyses confirmed that occurrences of more than 7 coincidences were rare in the population recorded by the intracortical probe and were totally lacking in a substantial fraction of the trials (Fig 3G). Fig 3H shows the prediction for epicortical peri-stimulus-time histograms with inclusion of all spikes recorded at the intracortical probe (black traces) and for synchronized spikes (red traces) with increasing numbers of synchronized units. Only for the highest number of coincidences, the peri-stimulus-time histogram becomes similar to the response actually observed at the cortical surface (green traces).

**Fig 3.**
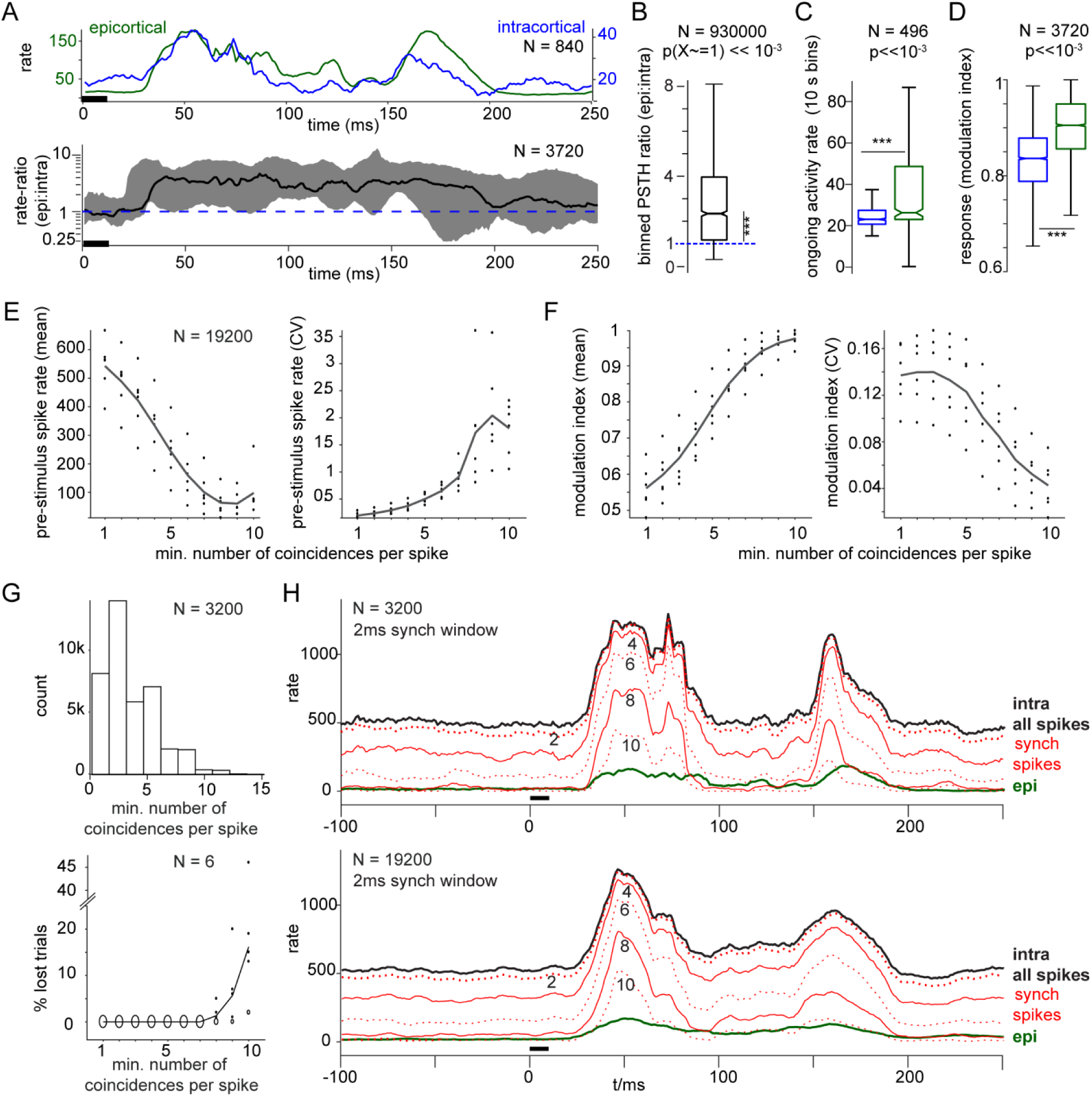
Relation between intracortical and epicortical spiking: ongoing activity and response dynamics. (A-D) Experimental data; (E-G) modeling; data in this figure were obtained from 3 animals; visual stimuli consisted of 10ms full-screen flashes of bright white light. (A) Trial-averaged PSTHs. (Top) Example of how trial-averaged epicortical responses (green) typically compare to the best-correlated intracortical response (blue); black horizontal bar, stimulus duration. (Bottom) Grand mean of the response ratio (epicortical over intracortical) over animals and trials. 1ms bins; shading: range of the data (minimum to maximum). (B) Data from A, collapsed across time bins and tested against a mean value of one, i.e., against equality of response strengths (N, number of trials; p-value: two-sided sign test against unity mean). (C) Firing-rate distributions of ongoing activity (10 s bins; N, number of bins; blue: intracortical; green: epicortical data). (D) Spike-rate modulation indices for intracortical (blue) and epicortical (green) responses to visual stimuli. (E-G) Forward modeling of epicortical signals based on the intracortical data from the same animals as in A-D; the forward model assumes a 2ms synchronicity window width. (E) Mean values (left) and coefficient of variation (CV, right) of pre-stimulus spiking as a function of the number of correlated intracortical spikes in the 2ms synchronicity window. N, number of trials. (F) Same statistics for relative response strength. (G) (Top) Distribution of bins with the required minimum number of coincident spikes. (Bottom) Percentage of lost´ trials that completely lacked bins with the required minimum number of coincidences. (H) Peri-stimulus-time histograms (10 ms boxcar-kernel convoluted spike trains) obtained by pooling over all 32 intracortical electrodes (black, intra all spikes), synchronous spikes (red, synch spikes) or epicortical spikes (green, epi). Italic numbers indicate the respective minimum number of coincidences per bin. Horizontal black bar: stimulus duration. (Top) Example data. (Bottom) same plot for grand averages across the three animals. Note that only the peri-stimulus-time histograms obtained with the highest number of coincidences resemble the epicortical response histograms.

### Synchronized spike signals map to the surface in a spatially specific manner

We conclude from the data so far that the epicortical multiunit signal may primarily reflect synchronized firing in the subjacent cortical columns. If so, the strength of correlations between intra- and epicortical spiking should decrease steeply with increasing spatial distance of the epicortical recording site to the intracortical probe. The steepness of this decay should be higher if only synchronized spikes are taken into account compared to the ‘all spikes’ condition. As shown in Fig 4A-C, these assumptions were confirmed for correlations between spike-density time series obtained for summed activity vs. synchronized spikes (1ms synchronicity window; 1ms boxcar kernel to produce spike-density series). For both ‘all spikes’ and ‘synch spikes’, correlation strength decays significantly (p<<0.001) with increasing distance to the intracortical probe. Still, the decay is substantially steeper for the synchronized spikes (exponential coefficients: -0.09 vs. -0.023), reflecting the more strongly localized peak in correlation between intra- and epicortical activity. In the example data set shown in Fig 4A-B, two correlation maxima in the ‘all spikes’ condition were located at larger distance from the intracortical probe. In contrast, a distinct hotspot neighboured the position of the intracortical probe in the ‘synch spikes’ condition. Thus, synchronized intracortical spikes are picked up mainly by nearby epicortical contacts.

To assess how different units along the shank of the intracortical probe contribute to the epicortical signal, we plotted the average depth-profile of best pair correlation strength along with the corresponding spike rate profiles (Fig 4D,E). Note that correlations between intra- and epicortical activities remain significant down to the deepest electrode contacts for both summed and synchronized spikes. However, a significant difference between the two conditions was only observed for the superficial populations. The profiles of spike rates (Fig 4E) confirm that this effect is not related to the depth distribution of spiking densities.

**Fig 4.**
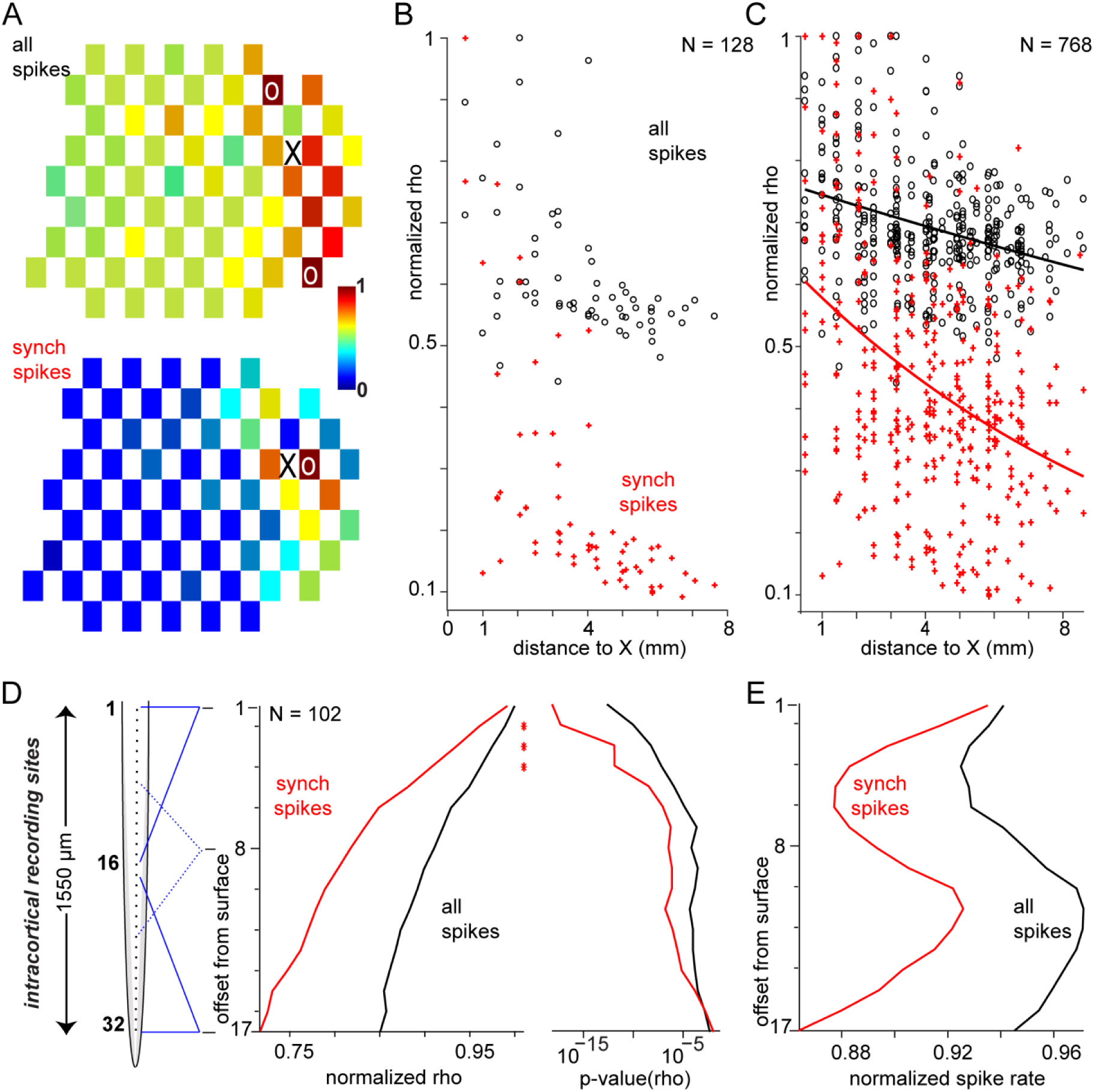
Surface topographies and depth profiles of correlations between intra-and epicortical spike trains. (A) Example topography of the strength of correlation (normalized to maximum) between intra- and epicortical spiking at different electrodes of the ECoG array. O, contact with best correlation; X, position of the intracranial probe. (Top) Results for all spikes from the intracortical probe. (Bottom) Results for synchronized spikes only (1ms synchronicity window; minimum number of 7 coincidences per bin). Note that the topography for ‘all spikes’ shows correlation over larger distance and includes maxima at higher spatial separation from the intracortical probe. (B) Same data collapsed into a distribution of correlation of spiking activity (normalized rho) as a function of distance from the intracortical probe. Black, data for ‘all spikes’; red, data for ‘synch spikes’. (C) Population statistic across all animals. (D) Depth-resolved correlation of spiking activity with the epicortical recording (computed for contacts with the best average correlation). To obtain a depth-resolved distribution, spike trains containing ‘all spikes’ or ‘synch spikes’ were generated for subsets of 16 neighbouring contacts of the intracortical probe (minimum number of coincidences per bin set to 4), with the offset from cortical surface varying from 1 to 17 contact positions. Red asterisks indicate a significant difference between the depth profiles for ‘synch spikes’ (red) vs. ‘all spikes’ (black) for offsets between 2 and 4. (E) Depth-resolved distribution of spike rate for ‘synch spikes’ (red) vs. ‘all spikes’ (black), averaged across neighbouring electrodes as in D.

## Discussion

Designs of ECoG electrode arrays are constantly being diversified and refined. In particular, designs that allow recordings of spiking activity are of increasing interest since they combine the technical advantages of ECoG arrays (such as flexibility, stability, lower invasiveness and large-scale coverage) and the information gained from measuring neural suprathreshold activity. However, the origin and physiological significance of epicortically measured multiunit spiking are still unexplored. Our study aimed to record multiunit activity during ongoing activity and sensory stimulation with a custom ECoG array of midsize (250 µm diameter) electrodes and to elucidate the relation of this epicortical spiking activity to its intracortical sources. We provide empirical and modeling data strongly suggesting that the epicortical electrodes used in our study selectively capture synchronized spiking of neurons in the subjacent cortex. As a result, responses to sensory stimulation are more robust and less noisy as compared to intracortical activity, and receptive field properties are well preserved in our epicortical recordings.

We show that the detection of spiking activity in epicortically recorded traces requires no modification of parameters such as frequency range or detection threshold. Remarkably, epicortical spike waveforms resembled intracortical spikes in shape, despite the presumed summation of intracortical spikes at near-zero lag which may give rise to their detectability at the cortical surface. Here, one may conceive modeling approaches to decipher whether the bimodality of peak-to-valley durations in epicortical spikes could result from two modes in the distribution of spike-spike lags that determine the outcomes of spike summation, rather than from signals reflecting contributions of fast vs. slow spikes.

Analyses of correlation patterns between spike-density time series suggest that the activity captured by a single epicortical contact conveys information of spiking units across the entire cortical depth, whereby the contribution is most prominent for the superficial layers. Considering the diameter of our ECoG electrodes (250 µm), it seems straightforward to assume that epicortical multiunit activity reflects signals from several neighbouring cortical columns. Overall, our data suggest that most of the signal propagated to the surface reflects synchronized spikes in the subjacent cortex. This may lead to a reduction of variability of responses and an increase in signal-to-noise ratio. Supporting this assumption, epicortical responses were higher in relation to ongoing activity and less variable than intracortical responses in terms of peak latency, amplitude and half-maximum duration. Importantly, the spike rates in ongoing activity and responses were not as high as expected for linear summation of spiking activity over dozens or hundreds of neural units but, rather, best predicted by taking only synchronized intracortical spikes into account.

The preservation of stimulus features in the epicortical signal was tested in both the visual and auditory domain. We found no significant differences between the bandwidth of auditory tuning curves (pure tone frequency) between ECoG and intracortical recordings. This seems surprising, as the ECoG electrode is likely to pool signals over neighbouring tonotopic columns. Possibly, the change in preferred frequency along the tonotopic axis [20] is still negligible compared to the diameter of the ECoG electrodes used here. As a result, the epicortical signal is reflecting underlying local activities which are still similar to each other, making the resultant sum a robust, yet stimulus specific signal. An analogue rationale may explain our observation that epicortically measured direction selectivity matches the values in the laminar recorded data.

Comparisons of visual receptive field sizes yielded a particular informative insight with respect to the origin of the epicortical multiunit signal. Across the population, receptive fields were smaller in diameter than those obtained from intracortical electrodes. Modeling of our data suggests that this effect can be predicted if only precisely synchronized spikes are taken into account, with an optimal synchrony window of ∼2 ms. With respect to receptive field width, we argue that optimal stimulation, i.e., confined illumination of the field’s hot spot, produces high firing rates and thus a high rate of spike-time coincidences. By contrast, responses to illumination in the periphery are expected to be much weaker and more variable in timing, hence producing substantially less coincidences. As a result, the flanks of the retinotopic tuning curve are damped and half-maximum width is decreased. Forward modeling of epicortical responses showed that they could be predicted best if increasing numbers of synchronized intracortical spikes were included.

Our data suggest that spikes become detectable at the cortical surface only if signals from a sufficiently high number of cortical cells are precisely synchronized. It seems likely that only precisely synchronised discharges can superimpose to give rise to epicortical signals that are large enough to be distinguished from noise and get detected as spike waveforms at the surface. This assumption is in line with previous experimental and modeling work which has suggested that highly synchronized intracortical spikes can even be detected in EEG recordings from the scalp [21,22,23].

One possibility is that the ECoG spikes result from summation of action potentials backpropagating into the apical dendrites of pyramidal cells. As shown by microelectrode recordings [24,25], such backpropagating action potentials generate significant extracellular fields. If strongly synchronized across the dendrites of neighbouring cells, this might give rise to fast spikes detectable at the cortical surface. Additional contributions might result from spikes originating in the dendrites which can be boosted by coincidence of inputs to apical dendrites with backpropagating action potentials [26], which is also highly likely to result from synchronized population activity. There is evidence to suggest that such dendritic spikes are detectable at the cortical surface [27].

## Conclusions

In summary, our results suggest that ECoG arrays can be used for selective recording of synchronized spikes, yielding robust responses with less jitter in latency, amplitude and duration as compared to single-unit or sparse multi-unit data. This reduction of noisy components that vary strongly across trials or between units allows single-trial based high-speed processing required for efficient training of response-decoders and real-time applications such as prosthetic device control or BMI-based communication [28]. Furthermore, the mesoscale approach pursued here may be particularly advantageous for characterization of large-scale networks by minimally invasive recordings in chronically implanted animals or even humans.

Combining the advantage of access to spike signals with large coverage, this ECoG-based approach may promote novel insights into large-scale network dynamics in behaving subjects. Synchronous spiking in particular has been proposed as a mechanism in the propagation of neural information, attentional gain modulation and solutions to the binding problem, among others [29,30,31,32,33]. Given the high relevance of neural synchrony for sensorimotor and cognitive processing, chronic implants that capture synchronized spiking of neural assemblies at the cortical surface are likely to increase the potential for decoding of signals in BMI approaches.

## Acknowledgements

We would like to thank Iain Stitt and Dorrit Bystron for assistance during surgery as well as Guido Nolte for comments on data analysis. This research was supported by funding from the German Research Foundation (SFB 936/A2, SPP 1665/EN533/13-1, SPP 2041/EN533/15-1, A.K.E.).

